# Slit-independent guidance of longitudinal axons by *Drosophila* Robo3

**DOI:** 10.1101/2023.05.08.539901

**Authors:** Abigail Carranza, LaFreda J. Howard, Haley E. Brown, Ayawovi Selom Ametepe, Timothy A. Evans

## Abstract

*Drosophila* Robo3 is a member of the evolutionarily conserved Roundabout (Robo) receptor family and one of three *Drosophila* Robo paralogs. During embryonic ventral nerve cord development, Robo3 does not participate in canonical Slit-dependent midline repulsion, but instead regulates the formation of longitudinal axon pathways at specific positions along the medial-lateral axis. Longitudinal axon guidance by Robo3 is hypothesized to be Slit dependent, but this has not been directly tested. Here we create a series of Robo3 variants in which the N-terminal Ig1 domain is deleted or modified, in order to characterize the functional importance of Ig1 and Slit binding for Robo3’s axon guidance activity. We show that Robo3 requires its Ig1 domain for interaction with Slit and for proper axonal localization in embryonic neurons, but deleting Ig1 from Robo3 only partially disrupts longitudinal pathway formation. Robo3 variants with modified Ig1 domains that cannot bind Slit retain proper localization and fully rescue longitudinal axon guidance. Our results indicate that Robo3 guides longitudinal axons independently of Slit, and that sequences both within and outside of Ig1 contribute to this Slit-independent activity.

## Introduction

Transmembrane receptor proteins in the Roundabout (Robo) family regulate a number of axon guidance decisions in bilaterian animals (Blockus and Chedotal, 2016; Dickson and Gilestro, 2006). The canonical role of Robo receptors is to regulate midline crossing of axons in the central nervous system (CNS) by signaling repulsion in response to their cognate Slit ligands, which are typically expressed at the CNS midline (Brose et al., 1999; Kidd et al., 1998; Kidd et al., 1999; Long et al., 2004; Zallen et al., 1998). In some animal groups, including vertebrates and insects, Robo family members have acquired other axon guidance roles in the CNS in addition to midline repulsion (Evans and Bashaw, 2010; Evans and Bashaw, 2012; Farmer et al., 2008; Jaworski et al., 2010; Jaworski et al., 2015; Long et al., 2004; Rajagopalan et al., 2000b; Sabatier et al., 2004; Simpson et al., 2000b). In *Drosophila*, three Robo receptors are present (Robo1, Robo2, and Robo3) and each plays a specialized set of roles during development of the embryonic CNS. Robo1 and Robo2 cooperate to signal midline repulsion in response to Slit, while Robo2 and Robo3 determine the medial-lateral position of longitudinal axon tracts within the neuropile of the ventral nerve cord (Rajagopalan et al., 2000a; Rajagopalan et al., 2000b; Simpson et al., 2000a; Simpson et al., 2000b). The differential roles of the three Robos in embryonic CNS development depend both on their distinct expression patterns and on structural differences between the receptors themselves (Evans and Bashaw, 2010; Spitzweck et al., 2010).

The overlapping expression patterns of Robo1, Robo2, and Robo3 in the developing embryonic ventral nerve cord reflect their differential roles in regulating the positions of axon pathways within the three dimensional structure of the nerve cord neuropile (Rajagopalan et al., 2000b; Simpson et al., 2000b). Robo1 protein is detectable on axons across the entire width of the longitudinal connectives and excluded from commissural axon segments, reflecting its broad role as the main midline repulsive Slit receptor (Kidd et al., 1998; Kidd et al., 1999). In *robo1* mutants, many axons ectopically cross the midline due to reduced midline repulsion. Robo2 is also broadly expressed in the early stages of axon guidance, where it cooperates with Robo1 to signal Slit-dependent repulsion, but becomes more restricted at later stages when it controls the formation of longitudinal pathways in the lateral-most region of the neuropile (Rajagopalan et al., 2000a; Rajagopalan et al., 2000b; Simpson et al., 2000a; Simpson et al., 2000b). Accordingly, *robo2* mutant embryos display mild ectopic midline crossing combined with defects in lateral axon pathway formation. Robo3 appears to play no role in regulating midline crossing in the embryonic nerve cord, but instead regulates longitudinal pathway formation in the intermediate region of the neuropile. Robo3 protein is restricted to the lateral two-thirds of the axon scaffold (intermediate plus lateral), and intermediate axon pathways are defective in *Robo3* mutant embryos (Rajagopalan et al., 2000b; Simpson et al., 2000b).

Robo1, Robo2, and Robo3 share an evolutionarily conserved ectodomain structure combining five immunoglobulin-like (Ig) domains with three fibronectin type-III (Fn) repeats (Kidd et al., 1998; Rajagopalan et al., 2000b; Simpson et al., 2000a). Although all three *Drosophila* Robos can act as Slit receptors (Brose et al., 1999; Rajagopalan et al., 2000b), we don’t yet have a complete understanding of which Robo-dependent axon guidance decisions are Slit-dependent, and which are Slit-independent. We have previously shown that deleting the N-terminal Ig1 domain from Robo1 abolishes Slit binding and also disrupts its ability to regulate midline repulsion in embryonic neurons (Brown et al., 2015). Similarly, deleting Ig1 from Robo2 prevents it from binding Slit and also disrupts its ability to signal midline repulsion and regulate lateral pathway formation (Evans et al., 2015; Howard et al., 2021). The Ig1 domains of Robo1 and Robo2 appear to have differential roles in protein localization in vivo, as deleting the Ig1 domain from Robo1 has no effect on its localization in embryonic neurons, but deleting Ig1 from Robo2 alters its distribution in neurons. Compared to full-length Robo2, Robo2ΔIg1 protein is present at lower levels on neuronal axons and correspondingly higher levels in neuronal cell bodies (Brown et al., 2015; Howard et al., 2021). It is not yet clear whether Ig1-dependent localization of Robo2 is separable from its ability to bind Slit, and whether the reduced axon guidance function of Robo2ΔIg1 is due solely to loss of Slit binding, its altered localization, or both (Howard et al., 2021).

Here, we examine the functional importance of Robo3 Ig1 for Slit binding, axonal localization of Robo3 protein in embryonic neurons, and guidance of longitudinal axons in the ventral nerve cord. We find that, similar to Robo1 and Robo2, deleting the Ig1 domain from Robo3 abolishes Slit binding, supporting the model that all three paralogs interact with Slit via a conserved Ig1 interface. Robo3ΔIg1 protein exhibits altered localization in embryonic neurons, similar to Robo2ΔIg1, yet still partially rescues Robo3’s role in guiding intermediate longitudinal axons, indicating that one or more additional Robo3 sequences also contribute to axon guidance independently of Ig1. Using a series of modified Robo3 Ig1 variants we show that the Slit binding, protein localization, and axon guidance functions of Robo3 Ig1 are genetically separable. We provide evidence that Robo3’s restricted expression pattern in the embryonic CNS is a result of a cell-autonomous function of Robo3 in excluding Robo3-expressing axons from the medial zone, suggesting mechanistic differences in the guidance of intermediate and lateral longitudinal axons by Robo3 and Robo2, respectively. Finally, we show that Robo3 variants with reduced or absent Slit binding can fully rescue its role in intermediate axon pathway formation. Together, this indicates that Robo3 guides intermediate axons in the *Drosophila* embryonic ventral nerve cord independently of Slit.

## Results

### The Ig1 domain of Robo3 is required for Slit binding

All three Robo receptors in *Drosophila* (Robo1, Robo2, and Robo3) share a common ligand, Slit (Brose et al., 1999; Rajagopalan et al., 2000b). For Robo1 and Robo2, it has been established that the first immunoglobulin-like domain (Ig1) is necessary for Slit binding (Brown et al., 2015; Evans et al., 2015; Howard et al., 2021). In Robo1, each of the other four Ig domains (Ig2-Ig5) are dispensable for Slit binding, and Robo2’s Ig2 domain, at least, is also dispensable for Slit binding (Brown and Evans, 2020; Evans et al., 2015; Reichert et al., 2016).

To test whether the Ig1 domain of Robo3 is similarly required for Slit binding, we expressed a variant of Robo3 from which Ig1 has been deleted (Robo3ΔIg1) in cultured *Drosophila* cells and examined its ability to bind Slit compared to full-length Robo3. We found that Slit bound strongly to *Drosophila* S2R+ cells expressing full-length Robo3, but not to untransfected cells or cells expressing Robo3ΔIg1 **(Figure 1A-C)**. N-terminal HA epitope tags on both variants revealed that Robo3 and Robo3ΔIg1 were expressed at similar levels and present at the surface of transfected cells. Thus, like *Drosophila* Robo1 and Robo2, the Ig1 domain of Robo3 is required for interaction with Slit.

**Figure 1.**
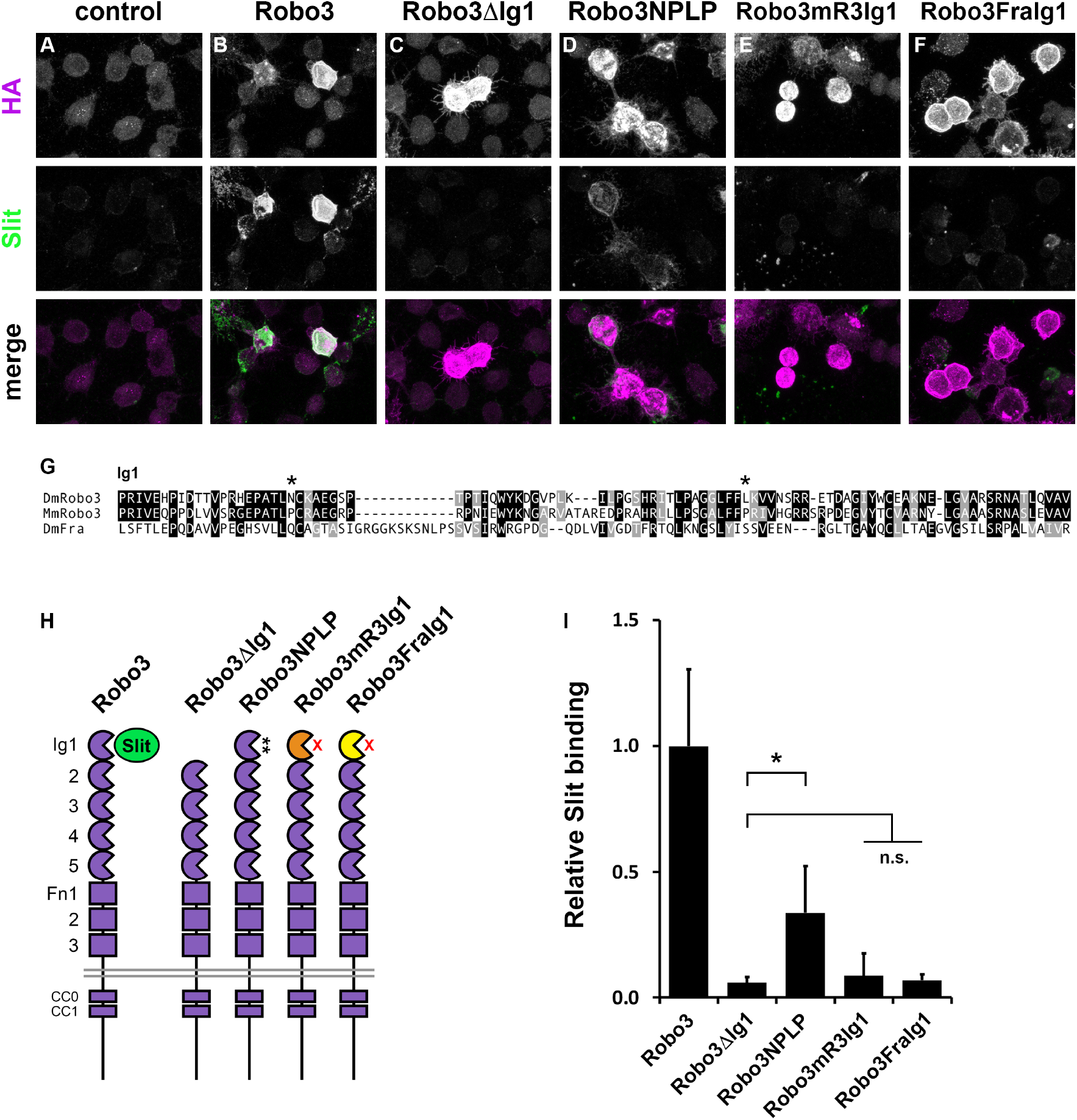
The Robo3 Ig1 domain is required for Slit binding. **(A-F)**Cultured *Drosophila* S2R+ cells were transfected with the indicated HA-tagged *UAS-Robo3* transgenes, then treated with Slit-conditioned media. After Slit treatment, cells were fixed and stained with anti-HA antibodies to detect HA-tagged Robo3 variants (magenta) and anti-Slit antibodies to detect bound Slit (green). Slit binds robustly to cells expressing full-length Robo3 **(B)**, but not untransfected cells **(A)**or cells expressing Robo3ΔIg1 **(C)**, Robo3mR3Ig1 **(E)**, or Robo3FraIg1 **(F)**variants. Slit binds weakly to cells expressing Robo3NPLP **(D). (G)**Amino acid alignment of Ig1 domains from *Drosophila* Robo3 (DmRobo3), mouse Robo3 (MmRobo3) and *Drosophila* Frazzled (DmFra). Identical amino acids are shaded black; similar amino acids are shaded gray. Asterisks indicate positions of N43P and L82P point mutations introduced into *Drosophila* Robo3 Ig1 to create the Robo3NPLP variant (Zelina et al., 2014). **(H)**Schematic of Robo3 variants tested for Slit binding. Robo3ΔIg1 has the Ig1 domain deleted; Robo3NPLP carries two point mutations in Ig1 (N43P and L82P); Robo3mR3Ig1 has the Ig1 domain replaced with Ig1 from mouse Robo3; Robo3FraIg1 has the Ig1 domain replaced with Ig1 from *Drosophila* Frazzled. **(I)**Bar graph quantifies Slit binding levels in conditions **B-F**, normalized to the level observed with full-length Robo3 **(B)**. Error bars show s.d. Robo3NPLP exhibited detectable levels of Slit binding (p<0.0001 compared to Robo3ΔIg1 by Student’s two-tailed t-test, but Robo3mR3Ig1 and Robo3FraIg1 were not significantly different than Robo3ΔIg1 (p>0.1 for each).

### No trans-heterozygous interaction between *Robo3* and *slit*

As a first step in addressing whether *Robo3* and *slit* cooperate to guide longitudinal axons, we looked for a trans-heterozygous interaction between *Robo3* and *slit* in FasII-positive intermediate longitudinal axons. In wild-type embryos, FasII-positive axon pathways form in three distinct medial-lateral zones within the neuropile (medial, intermediate, and lateral). *Robo3* is required for the formation of FasII-positive pathways of the intermediate zone. In *Robo3* mutant embryos, these pathways fail to form properly, and instead, intermediate longitudinal axons shift closer to the midline to merge with pathways in the medial zone (Rajagopalan et al., 2000b). The initial models of Robo3’s function in guiding longitudinal axons posited that all three Robo receptors (Robo1, Robo2, and Robo3) act as repulsive Slit receptors in order to guide FasII-positive longitudinal axons to appropriate medial-lateral positions (Rajagopalan et al., 2000b; Simpson et al., 2000b). These models hypothesized that axons expressing Robo1+Robo3 select intermediate positions within the neuropile because they are repelled away from the midline more strongly than Robo1-expressing axons in the medial zone, but not as strongly as axons in the lateral zone expressing all three Robos. In *slit* mutant embryos, all of the longitudinal pathways collapse at the midline due to a complete loss of midline repulsion (Kidd et al., 1999; Rothberg et al., 1990). This strong midline collapse phenotype prevents any examination of longitudinal pathway formation, making it difficult to discern which other *robo-* dependent axon guidance activities may be deficient in *slit* mutants.

Zlatic et al (2003) reported that *Robo3* and *slit* function together to target chorodotonal sensory axon terminals to the intermediate zone, as chorodotonal projections displayed significant targeting defects in the ventral nerve cords of animals heterozygous for both *Robo3* and *slit*, but not *slit/+* or *Robo3/+* single heterozygotes (Zlatic et al., 2003). However, FasII-positive pathways appeared normal in these double heterozygous embryos. Consistent with this, we also observed no significant defects in intermediate FasII pathways in embryos heterozygous for strong loss of function mutations in both *slit* and *Robo3*, similar to animals that are heterozygous for either gene alone **(Figure 2A-C)**.

**Figure 2.**
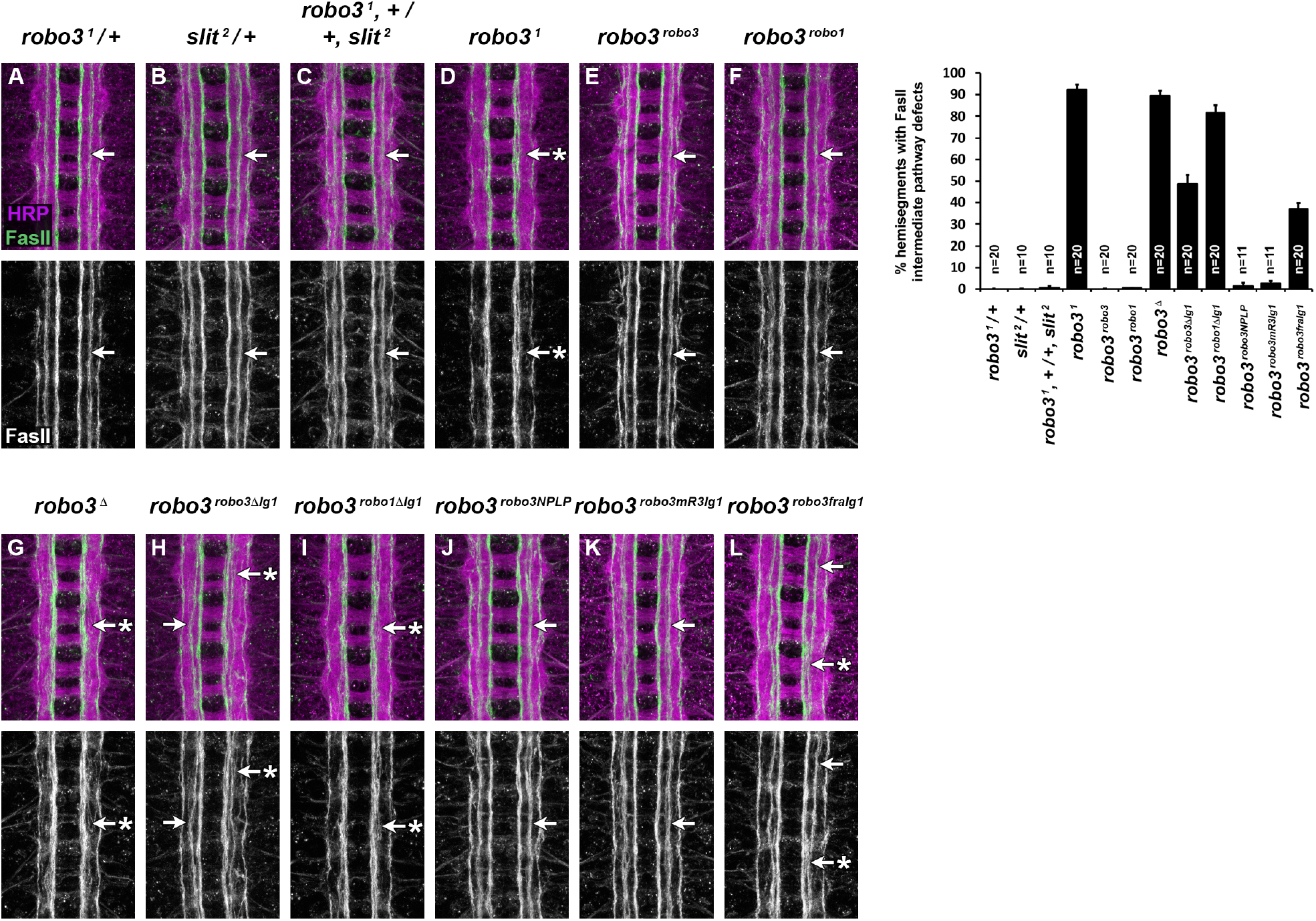
FasII-positive longitudinal pathway formation in *Robo3* modified embryos. **(A-L)**Stage 16-17 *Drosophila* embryonic ventral nerve cords stained with anti-HRP (magenta; labels all axons) and anti-FasII (green; labels a subset of longitudinal axon pathways) antibodies. Lower images show anti-FasII channel alone from the same embryos. **(A)**In *Robo3* ^*1*^*/+* heterozygous embryos, the wild type pattern of FasII-positive longitudinal axon pathways in three discrete zones (medial, intermediate, and lateral) is visible. The arrow denotes the intermediate pathway. **(B**,**C)**The intermediate FasII pathway also forms correctly in heterozygous *slit*^*2*^*/+* embryos **(B, arrow)**and in embryos heterozygous for both *slit* and *Robo3***(C, arrow)**. In homozygous *Robo3*^*1*^mutants, FasII-positive axons that normally form the intermediate pathway are displaced medially, and the intermediate pathway fails to form **(D, arrow with asterisk)**. In embryos where *Robo3* is replaced by a full-length *Robo3*cDNA **(E)**or a full-length *robo1* cDNA **(F)**, the intermediate pathway again forms correctly **(E**,**F, arrows). (G)** Embryos homozygous for a *Robo3* null deletion allele (*Robo3*^*Δ*^) exhibit intermediate pathway defects similar to *Robo3*^*1*^ embryos **(arrow with asterisk). (H)**In embryos homozygous for the *Robo3*^*Robo3ΔIg1*^allele, the intermediate pathway fails to form in around half of segments **(arrow with asterisk)**but forms relatively normally in the other half **(arrow). (I)**In embryos homozygous for the *Robo3*^*robo1ΔIg1*^allele, the intermediate pathway is defective in nearly all segments **(arrow with asterisk)**. The intermediate pathway forms normally in embryos homozygous for the *Robo3*^*Robo3NPLP*^**(J)**or *Robo3*^*Robo3mR3Ig1*^**(K)**alleles. In embryos homozygous for the *Robo3*^*Robo3FraIg1*^allele **(L)**, the intermediate FasII pathway forms normally in nearly two-thirds of segments **(arrow)**and is defective in the remaining segments **(arrow with asterisk)**. Bar graph shows quantification of intermediate FasII pathway defects in the genotypes shown in **(A-L)**. Error bars indicate standard error of the mean. Number of embryos scored for each genotype (n) is indicated.

Although trans-heterozygous interactions are often interpreted as evidence that two genes function together in the same genetic pathway, the absence of such an interaction does not demonstrate that the two genes do not function together. To definitively test whether Robo3 guides longitudinal axons in response to Slit, we set out to examine the *in vivo* axon guidance activity of our Slit-binding deficient Robo3ΔIg1 variant by expressing it in developing embryos in place of endogenous Robo3. We have used similar structure-function approaches to examine Slit-dependent activities of Robo1 and Robo2 (Brown et al., 2015; Evans et al., 2015; Howard et al., 2021).

### CRISPR/Cas9-based gene replacement of *Robo3*

To examine whether Slit binding is necessary for the axon guidance activity of Robo3 *in vivo*, we used a CRISPR/Cas9-based gene modification approach (Gratz et al., 2014; Port et al., 2014) to replace exons 2-12 of the *Robo3* gene with an HA-tagged *Robo3ΔIg1* cDNA **(Figure 3)**. In this modified allele, the endogenous *Robo3* promoter, transcriptional start site, first exon (including the start codon and signal sequence), and 16.6 kb first intron remain unmodified. Spitzweck et al. (2010) used a knock-in approach to similarly replace *Robo3* with a full-length *Robo3* cDNA, and showed that HA-tagged Robo3 protein expressed from this modified locus was properly expressed and could fully rescue Robo3’s role in guidance of longitudinal axons (Spitzweck et al., 2010). We have previously used CRISPR to replace *Robo3* with its ortholog from the flour beetle *Tribolium castaneum, TcRobo2/3*, and showed that TcRobo2/3 can fully rescue *Drosophila* Robo3’s role in intermediate pathway formation (Evans, 2017). Here, we used the same two guide RNAs (gRNAs) targeting exon 2 and exon 12, and the same homologous donor plasmid backbone, in this case containing the *Robo3ΔIg1* coding sequence.

**Figure 3.**
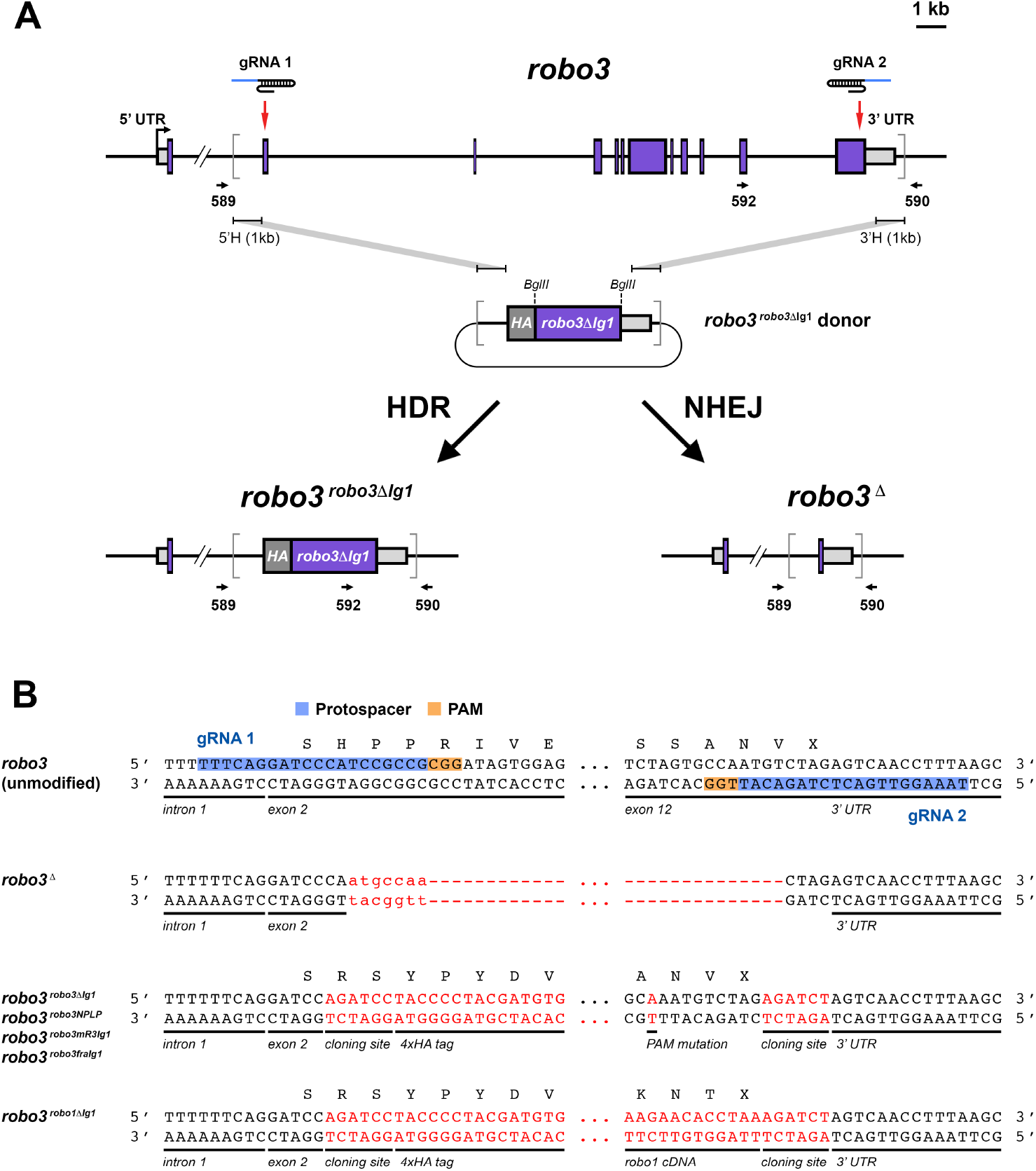
CRISPR-based gene replacement of *Robo3*. **(A)**Schematic of the *Robo3* gene showing intron/exon structure and location of gRNA target sites, *Robo3*^*Robo3ΔIg1*^homologous donor plasmid, and the modified alleles produced via homology directed repair (HDR) or non-homologous end joining (NHEJ). Endogenous *Robo3* coding exons are shown as purple boxes; 5’ and 3’ untranslated regions (UTR) are shown as light grey boxes. The start of transcription is indicated by the bent arrow. Introns and exons are shown to scale, with the exception of the first intron, from which approximately 13 kb has been omitted. Red arrows indicate the location of upstream (gRNA 1) and downstream (gRNA 2) gRNA target sites. Grey brackets demarcate the region to be replaced by sequences from the donor plasmid. Arrows under schematic indicate the position and orientation of PCR primers. **(B)**Partial DNA sequences of the unmodified *Robo3* gene and the modified alleles produced in this study. Black letters indicated endogenous DNA sequence; red letters indicate exogenous sequence. Both DNA strands are illustrated. The gRNA protospacer and PAM sequences are indicated for both gRNAs. In the *Robo3*^*Δ*^allele, all sequence between the two gRNA target sites (approximately 20 kb) is deleted, with 7 bp inserted at the deletion site. The first five base pairs of *Robo3* exon 2 are unaltered in the *Robo3*^*Robo3ΔIg1*^, *Robo3*^*Robo3NPLP*^, *Robo3*^*Robo3mR3Ig1*^, *Robo3*^*Robo3FraIg1*^, and *Robo3*^*robo1ΔIg1*^alleles; and the *Robo3* coding sequence beginning with codon H21 is replaced by the respective HA-tagged cDNA sequences. The endogenous *Robo3* transcription start site, ATG start codon, and signal peptide are retained in exon 1 of all modified alleles. The PAM sequences and portions of both protospacers are deleted or mutated in each HDR modified allele, ensuring that the HDR donor plasmids and final modified alleles are not targeted by Cas9. UTR, untranslated regions; 5’H, 5’ homology region; 3’H, 3’ homology region; HA, hemagglutinin epitope tag; gRNA, guide RNA; HDR, homology directed repair; PAM, protospacer adjacent motif.

For all CRISPR lines described here, donor and gRNA plasmids were co-injected into transgenic *Drosophila* embryos expressing Cas9 nuclease under control of the germline-specific *nanos* promoter (*nos-Cas9*.*P*) (Port et al., 2014), and F1 progeny from the injected flies were screened by PCR to identify those carrying the expected modified alleles. We generated stable lines derived from positive F1 flies and sequenced the modified locus fully from at least one line for each modified allele. Additional details are provided in the Methods and **Table 1**.

**Table 1.**
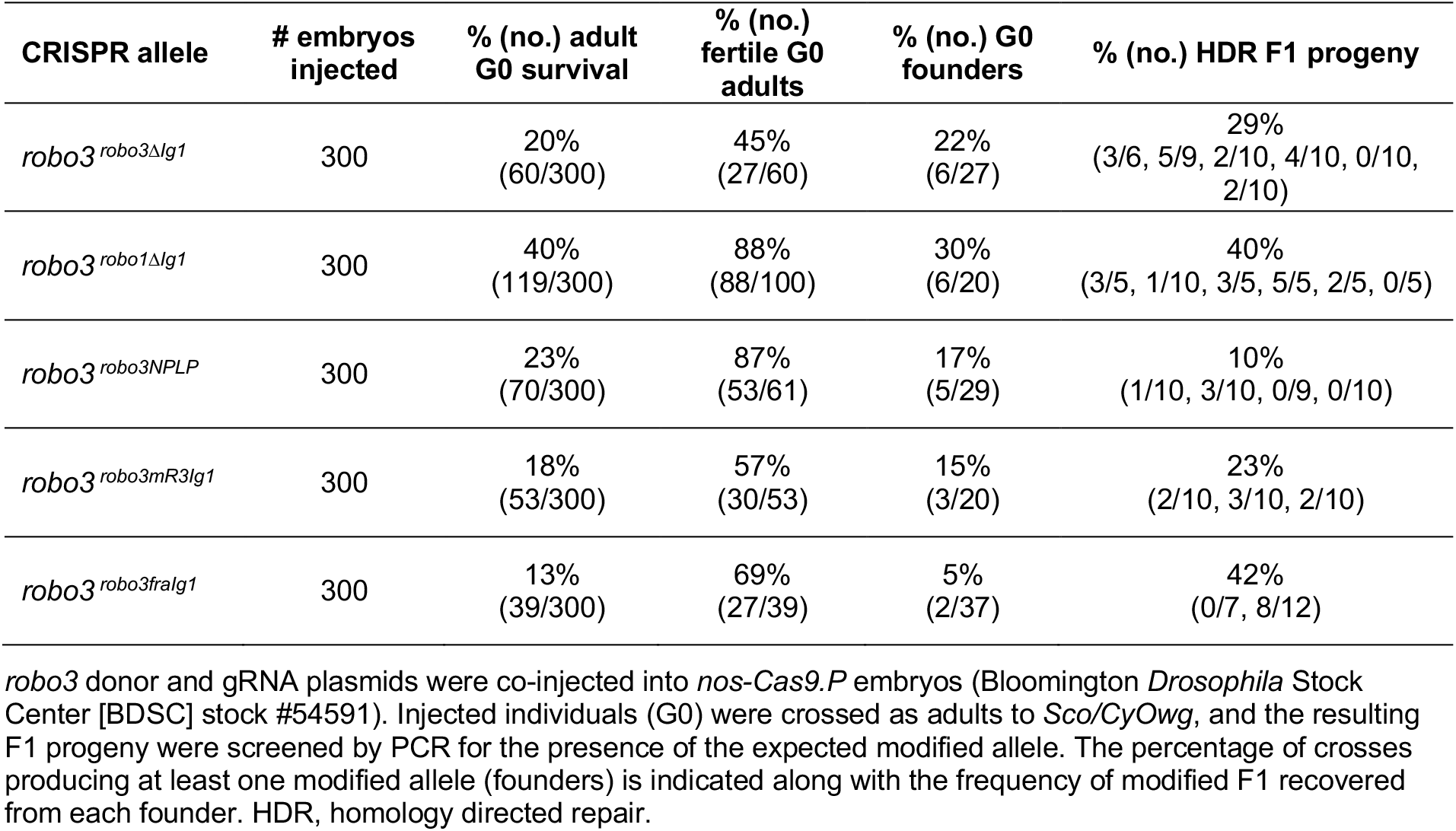
Survival, fertility, and transmission rates for CRISPR replacement of robo3.

### Recovery of an NHEJ-derived deletion of the *Robo3* locus

Our strategy of using two gRNAs to engineer *Robo3* via homology-directed repair (HDR) also allowed us to use PCR to screen for non-homologous end joining (NHEJ) events. We expected NHEJ repair to produce a deletion of all sequence between the two gRNA target sites, removing all but one coding exon and creating a null deletion allele of *Robo3*. Indeed, we detected one such deletion (designated *Robo3*^*Δ*^) in F1 flies derived from a G0 founder fly that also produced HDR progeny **(Figure 3)**. Thus, both NHEJ and HDR modifications can be recovered from the same injected fly when gRNA and donor plasmids are co-injected.

We used PCR primers flanking the gRNA target sites to amplify a PCR product containing the *Robo3*^*Δ*^deletion and sequenced this product to identify the deletion breakpoints. Consistent with previous descriptions of NHEJ repair in *Drosophila* (Bassett et al., 2013; Gratz et al., 2013; Kondo and Ueda, 2013; Yu et al., 2013), we found that the deletion breakpoints are located within the two protospacer sequences, 4-7 bp upstream of the PAM sequence for each gRNA target site. In addition to the 20,091 bp deleted in the *Robo3*^*Δ*^allele, 7 bp of unrelated sequence is inserted at the site of the deletion **(Figure 3B)**.

As the majority of coding exons are deleted in the *Robo3*^*Δ*^allele, we predicted that it would be a protein null allele, and that we would observe the same defects in longitudinal pathway formation that have previously been described for the strongly hypomorphic *Robo3*^*1*^point mutant allele (Evans, 2017; Rajagopalan et al., 2000b; Spitzweck et al., 2010). Indeed this was the case, as Robo3 protein was undetectable in the ventral nerve cord of homozygous *Robo3*^*Δ*^ embryos, similar to homozygous *Robo3*^*1*^ embryos **(Figure 4A-C)**, and intermediate axon pathway defects were present to a similar extent in *Robo3*^*1*^and *Robo3*^*Δ*^ embryos **(Figure 2D,G**).

**Figure 4.**
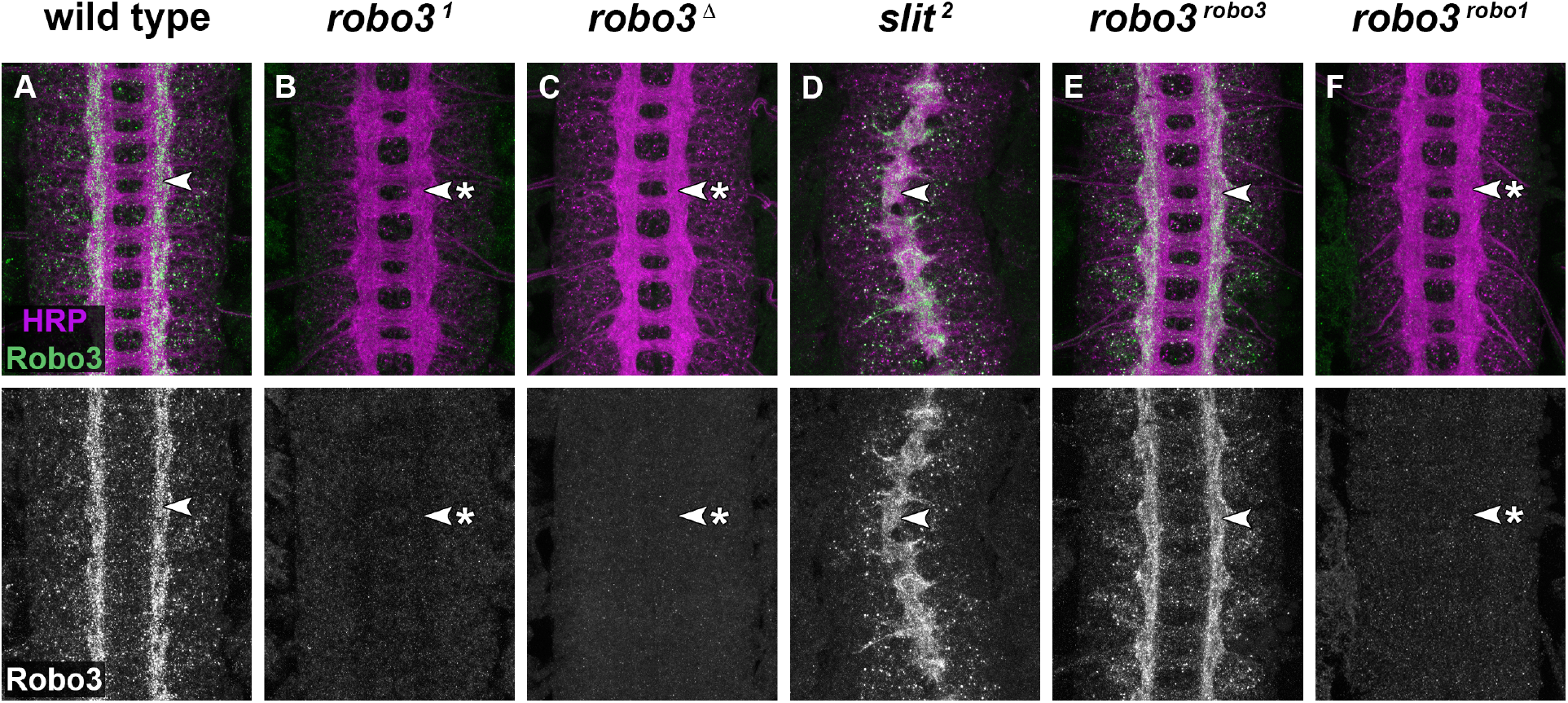
Endogenous Robo3 protein expression in wild type, mutant, and modified *Robo3*alleles. **(A-F)**Stage 16-17 *Drosophila* embryonic ventral nerve cords stained with anti-HRP (magenta; labels all axons) and anti-Robo3 (green) antibodies. Lower images show anti-Robo3 channel alone from the same embryos. In wild type embryos, endogenous Robo3 protein is detectable on longitudinal axons within the outer two-thirds of the neuropile **(A, arrowhead)**. Robo3 protein is undetectable in embryos homozygous for the loss of function *Robo3*^*1*^allele **(B, arrowhead with asterisk)**(Rajagopalan et al., 2000b) or the *Robo3*^*Δ*^deletion allele **(C, arrowhead with asterisk)**. There are no large-scale defects detectable with anti-HRP in the axon scaffold of homozygous *Robo3*^*1*^or *Robo3*^*Δ*^mutants. In homozygous *slit*^*2*^null mutants, all axons collapse at the midline due to a complete lack of midline repulsion (Kidd et al., 1999; Rothberg et al., 1990), but Robo3 protein is still localized properly to axons **(D, arrowhead)**. In embryos in which the *Robo3* gene has been replaced with an HA-tagged *Robo3* cDNA, Robo3 protein expressed from the modified locus reproduces its normal expression pattern **(E, arrowhead)**(Spitzweck et al., 2010), while Robo3 protein is absent from embryos in which the *Robo3* gene has been replaced with an HA-tagged *robo1* cDNA **(F, arrowhead with asterisk)**(Spitzweck et al., 2010).

### Expression and localization of Robo3ΔIg1 in embryonic neurons

To examine whether the Ig1 domain of Robo3 is required for its proper expression and localization in embryonic neurons, we used an antibody against the N-terminal HA tag in our *Robo3*^*Robo3ΔIg1*^allele to compare the expression of Robo3ΔIg1 protein with endogenous Robo3 in wild type embryos (detected with a monoclonal antibody raised against the Robo3 cytodomain) along with HA-tagged full-length Robo3 expressed from the *Robo3*^*Robo3*^ modified knock-in allele (Spitzweck et al., 2010) **(Figure 5)**. Robo3 protein is primarily localized to longitudinal axons in the lateral two-thirds of the neuropile in late-stage wild type embryos **(Figure 4A)**(Rajagopalan et al., 2000b; Simpson et al., 2000b), and this expression pattern is faithfully reproduced by anti-HA staining in *Robo3*^*Robo3*^ embryos **(Figure 5A)**(Spitzweck et al., 2010).

**Figure 5.**
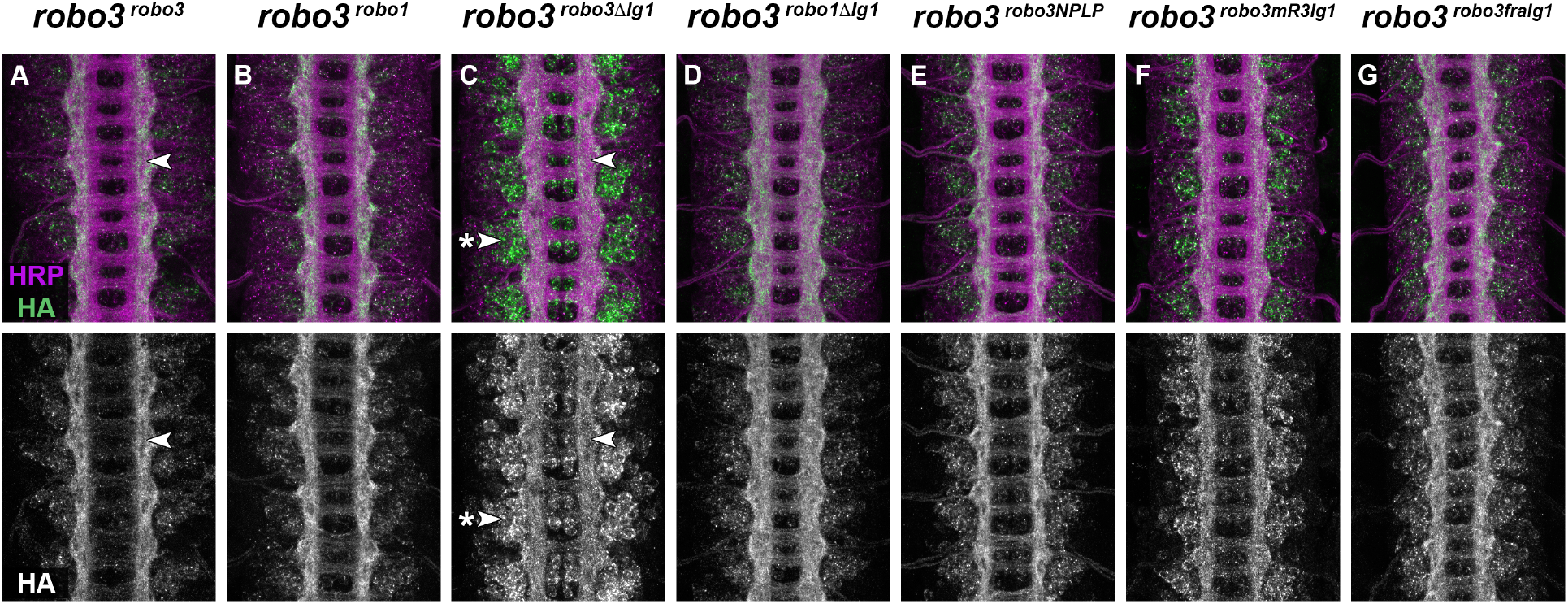
Expression of HA-tagged *Robo3* alleles in the embryonic CNS. **(A-G)**Stage 16-17 *Drosophila* embryonic ventral nerve cords stained with anti-HRP (magenta; labels all axons) and anti-HA (green) antibodies. Lower images show anti-HA channel alone from the same embryos. **(A)**In homozygous *Robo3*^*Robo3*^ embryos, HA-tagged Robo3 protein expressed from the modified locus reproduces Robo3’s endogenous expression pattern: it is localized to longitudinal axons and restricted to the intermediate and lateral zones **(A, arrowhead). (B)**Robo1 protein expressed from the *Robo3* locus in homozygous *Robo3*^*robo1*^ embryos shows equivalent axonal localization and exclusion from the medial zone. In homozygous *Robo3*^*Robo3ΔIg1*^ embryos, HA-tagged Robo3ΔIg1 protein is present on longitudinal axons but is not excluded from the medial zone **(C, arrowhead)**. Robo3ΔIg1 protein is also detectable at elevated levels in neuronal cell bodies within the cortex **(C, arrowhead with asterisk). (D)**In homozygous *Robo3*^*robo1ΔIg1*^ embryos, HA-tagged Robo1ΔIg1 protein is primarily localized to axons but not excluded from the medial zone. **(E-G)**HA-tagged Robo3NPLP, Robo3mR3Ig1, and Robo3FraIg1 proteins are primarily localized to axons in homozygous *Robo3*^*Robo3NPLP*^, *Robo3*^*Robo3mR3Ig1*^, and *Robo3*^*Robo3FraIg1*^ embryos, respectively. Robo3mR3Ig1 and Robo3FraIg1 show slightly elevated expression in cell bodies. Robo3NPLP and Robo3mR3Ig1 proteins are properly excluded from the medial zone **(E**,**F)**but Robo3FraIg1 protein is not **(G)**.

We have previously shown that deleting the Ig1 domain from *Drosophila* Robo1 does not affect its expression pattern, axonal localization, clearance from commissural axon segments, or regulation by the endosomal sorting protein Commissureless (Comm) (Brown et al., 2015), while deleting the Ig1 domain from Robo2 does appear to affect its proper localization to neuronal axons (Howard et al., 2021). In homozygous *Robo3*^*Robo3ΔIg1*^ embryos, we observed two deviations from the normal Robo3 expression pattern. First, while HA-tagged Robo3ΔIg1 protein was clearly detectable on axons, it was also present at elevated levels in neuronal cell bodies relative to full-length Robo3. Second, Robo3ΔIg1 protein was not restricted to axons in the lateral two-thirds of the neuropile, as Robo3 normally is, but rather was detectable across the entire width of the longitudinal axon scaffold **(Figure 5C)**. We have previously reported similarly elevated cell body expression of Robo2ΔIg1 in homozygous *robo2*^*robo2ΔIg1*^ embryos, although we did not see any medial expansion of Robo2ΔIg1 axonal expression like we do here for Robo3ΔIg1 (Howard et al., 2021).

One possible explanation for the mislocalization of Robo3ΔIg1 protein is that Slit binding may be required for proper localization of Robo3, perhaps by stabilizing Robo3 protein at the growth cone surface, preventing its endocytosis, or through some other mechanism. To test this, we examined endogenous Robo3 protein expression in *slit* null mutants using the anti-Robo3 antibody. We found that axonal localization of Robo3 does not depend on Slit, as endogenous Robo3 protein remains localized to neuronal axons in *slit* mutant embryos and does not appear to be elevated in neuronal cell bodies relative to wild type embryos **(Figure 4A,D**). This result suggests that the Robo3 Ig1 domain contributes to axonal localization of Robo3 protein in a Slit-independent manner.

### Robo3ΔIg1 protein is mislocalized in Robo1-expressing neurons

We reasoned that the differences in protein localization between Robo3 and Robo3ΔIg1 might be due to an artefact of the CRISPR modification, an effect on folding or stability of the Robo3ΔIg1 protein due to the deletion of Ig1, or a previously uncharacterized requirement for Ig1 for proper trafficking of Robo3 to neuronal axons. In addition, given that we did not observe any effects on localization of Robo1 in previous studies when we deleted its Ig1 domain (Robo1ΔIg1) (Brown et al., 2015), we wondered whether this might represent a difference between the Robo1 and Robo3 proteins, or instead a differential requirement for Ig1-dependent trafficking in neurons that normally express Robo3 versus neurons that normally express Robo1 (these populations should not be mutually exclusive, as both intermediate and lateral axons are predicted to co-express Robo1 and Robo3).

We found that the mislocalization of Robo3ΔIg1 compared to full-length Robo3 is not a unique feature of Robo3-expressing neurons or an artefact of CRISPR modification at the *Robo3* locus, as we observed a similar difference in localization when we expressed both versions of Robo3 in Robo1-expressing neurons using a *robo1* genomic rescue transgene (Brown et al., 2015) **(Figure S1)**. In these embryos, full-length Robo3 is detectable across the entire width of the longitudinal connectives, similar to Robo1’s normal expression pattern, while Robo3ΔIg1 is present at a lower level on axons and at elevated levels in neuronal cell bodies. In contrast, Robo1ΔIg1 protein expressed from the same rescue construct is primarily localized to axons, similar to full-length Robo1 and Robo3 (Brown et al., 2015) **(Figure S1)**. This suggests that the differential effect on localization of deleting Ig1 from Robo1 and Robo3 is not due to their expression in distinct neuronal subsets, but rather may reflect differential roles of the Ig1 domain in axonal localization of Robo1 vs Robo3.

### Robo1ΔIg1 protein is properly localized to axons in Robo3-expressing neurons

If the differential effect of Ig1 deletion on localization of Robo1ΔIg1 and Robo3ΔIg1 proteins is due to different requirements for the Ig1 domain in Robo1 and Robo3 protein localization, we would predict that Robo1ΔIg1 protein would exhibit proper axonal localization when expressed in Robo3’s normal expression pattern. To test this, we generated a CRISPR-modified allele in which the *Robo3* coding sequence is replaced with an HA-tagged *robo1ΔIg1* cDNA and examined the distribution of Robo1ΔIg1 protein in embryos carrying this allele using anti-HA. Indeed, we found that Robo1ΔIg1 protein is primarily localized to axons in *Robo3*^*robo1ΔIg1*^ embryos, similar to full-length Robo3 protein in wild type and *Robo3*^*Robo3*^ embryos, or full-length Robo1 protein in *Robo3*^*robo1*^ embryos **(Figure 5D)**. This demonstrates that Robo1ΔIg1 protein can be properly trafficked in the neurons that normally express Robo3, and therefore this trafficking is not dependent on Robo1’s Ig1 domain. This further suggests that the differences in protein localization we observe with Robo1ΔIg1 and Robo3ΔIg1 reflect a differential function of Robo1 and Robo3 Ig1 domains in protein trafficking, rather than differences in trafficking mechanisms between neuronal subsets (Robo1-expressing vs Robo3-expressing neurons).

### Robo3ΔIg1-expressing axons are misguided into the medial zone

We noted that while Robo1ΔIg1 protein is properly localized to axons in homozygous *Robo3*^*robo1ΔIg1*^ embryos, it is not restricted to axons in the lateral two-thirds of the neuropile. Instead, like Robo3ΔIg1 protein in *Robo3*^*Robo3ΔIg1*^ embryos, it is also detectable across the entire width of the axon scaffold **(Figure 5D)**. Thus, both Robo3ΔIg1 and Robo1ΔIg1 failed to reproduce Robo3’s normal restriction to intermediate and lateral regions. Since Robo3 is predicted to act cell-autonomously to direct axons to intermediate regions of the neuropile, we reasoned that the expansion of Robo3ΔIg1’s and Robo1ΔIg1’s expression domains to the medial zone might be caused by misguidance of normally Robo3-expressing axons into medial pathways.

To test this, we compared the distribution of Robo3ΔIg1 and Robo1ΔIg1 protein in homozygous *Robo3*^*Robo3ΔIg1*^and *Robo3*^*robo1ΔIg1*^ embryos with their heterozygous siblings (*Robo3*^*Robo3ΔIg1*^*/+*and *Robo3*^*robo1ΔIg1*^*/+)*, which carry one wild-type copy of *Robo3***(Figure 6)**. Indeed, we found that both Robo3ΔIg1 and Robo1ΔIg1 proteins are properly excluded from the medial zone in heterozygous *Robo3*^*Robo3ΔIg1*^*/+*and *Robo3*^*robo1ΔIg1*^*/+* embryos, respectively **(Figure 6C-F**). We infer that Robo3-expressing axons are guided inappropriately into the medial zone when they express Robo3ΔIg1 or Robo1ΔIg1 in place of Robo3, resulting in an expansion of their protein expression domains across the entire width of the longitudinal scaffold. These results are consistent with a cell-autonomous role for Robo3 in excluding Robo3-expressing axons from the medial zone of the embryonic ventral nerve cord, which can be fully rescued by expression of Robo1 (but not Robo3ΔIg1 or Robo1ΔIg1) in place of Robo3.

**Figure 6.**
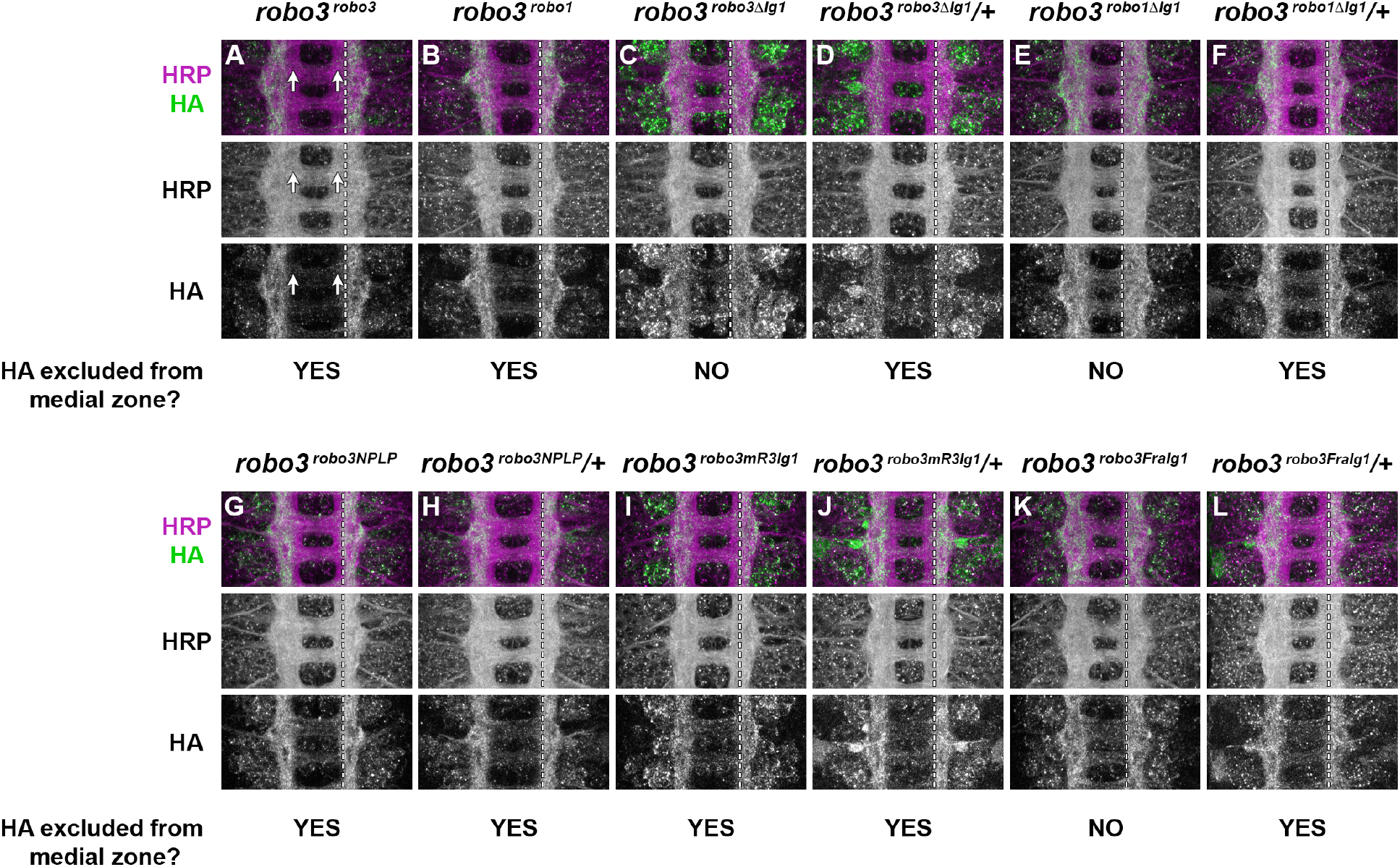
Comparison of protein localization in homozygous vs heterozygous HA-tagged *Robo3* alleles. **(A-L)**Single segments from stage 16-17 *Drosophila* embryonic ventral nerve cords stained with anti-HRP (magenta; labels all axons) and anti-HA (green) antibodies. Middle and lower images show anti-HRP and anti-HA channels alone from the same embryos, respectively. Dashed vertical lines show the medial boundary of the HA expression domain on the right half of the segment for each embryo. Homozygous *Robo3*^*Robo3*^ embryos **(A)**illustrate the normal exclusion of Robo3-expressing axons (labeled by anti-HA) from the medial zone of the neuropile **(A, arrows)**. HA-expressing axons are likewise excluded from the medial zone in homozygous *Robo3*^*robo1*^ embryos **(B)**. For the remaining genotypes, homozygous embryos are shown next to heterozygous sibling embryos. In all cases, the Robo1 and Robo3 variant proteins display similar levels of expression on axons and/or cell bodies in both homozygous and heterzygous siblings, and HA-expressing axons are excluded from the medial zone in heterozygous siblings even when they are not excluded from the medial zone in homozygous siblings, as with *Robo3*^*Robo3ΔIg1*^**(C**,**D)**, *Robo3*^*robo1ΔIg1*^**(E**,**F)**, and *Robo3*^*Robo3FraIg1*^**(K**,**L)**.

### Robo3ΔIg1 partially rescues intermediate pathway formation

To more closely examine longitudinal axon guidance in our modified *Robo3* embryos, and to determine whether Robo3ΔIg1 can substitute for Robo3 to promote the formation of specific longitudinal axon pathways in the intermediate zone, we used anti-FasII to examine intermediate pathway formation in embryos homozygous for our modified *Robo3*alleles. As controls, we included a set of *Robo3* knock-in alleles created by Spitzweck et al. (2010) in which the *Robo3* coding region is replaced with HA-tagged *Robo3* or *robo1* cDNAs. Intermediate FasII pathways form correctly in both of these backgrounds, confirming that removing most of the introns from *Robo3* and adding an N-terminal HA tag do not interfere with its function, and that full-length *robo1* can fully substitute for *Robo3* to promote intermediate pathway formation **(Figure 2E,F**)(Spitzweck et al., 2010).

We found that FasII-positive longitudinal pathways form properly in 51.4% of hemisegments in homozygous *Robo3*^*Robo3ΔIg1*^ embryos **(Figure 2H)**, demonstrating that Robo3ΔIg1 can partially substitute for Robo3 to guide longitudinal axons to the intermediate zone. Importantly, since Robo3ΔIg1 is partially mislocalized *in vivo*, we cannot distinguish whether its decreased activity is due strictly to the loss of an Ig1-dependent function (like Slit binding), or merely due to decreased levels of Robo3ΔIg1 protein on axons. In other words, Ig1 might play no direct role in Robo3-dependent guidance, and Robo3ΔIg1 might be fully functional if it were localized to axons at similar levels to full-length Robo3.

Interestingly, although intermediate FasII pathways form correctly when *Robo3*is replaced by full-length *robo1***(Figure 2F)**(Spitzweck et al., 2010), *Robo3*^*robo1ΔIg1*^ embryos fail to properly form intermediate axon pathways in 81.4% of hemisegments (Figure 2I). This result is puzzling as Robo1ΔIg1 protein is properly localized to axons **(Figure 5C)**. Thus, although Robo1ΔIg1 is properly localized in neurons that normally express Robo3, it cannot rescue intermediate pathway formation to the same extent as Robo3ΔIg1. This indicates that the Ig1-independent axon guidance activity we observe with Robo3ΔIg1 (which must be conferred by sequences outside of Ig1) is not shared by Robo1ΔIg1.

### Creation of Robo3 Ig1 variants that do not bind Slit

The above results suggest that Robo3’s Ig1 domain is required for its localization to axons, and since Robo3 protein is not mislocalized in *slit*mutants, this activity may be independent of its ability to bind Slit. We next attempted to separate Robo3 Ig1’s role in protein localization from its role in Slit binding by creating a version of Robo3 that retains an Ig1 domain but cannot bind Slit. Previous biochemical interaction and site-directed mutagenesis studies *in vitro*have identified evolutionarily conserved residues in the Ig1 domains of Robo receptors from insects and vertebrates that are necessary for Slit binding (Fukuhara et al., 2008; Morlot et al., 2007). Additionally, a more recent report identified naturally occurring amino acid substitutions in mammalian Robo3 Ig1 that prevent Slit binding (Zelina et al., 2014).

We made three separate modifications to *Drosophila* Robo3 to prevent Slit binding: first, we introduced two point mutations in Ig1 (N43P and L82P) to mimic two of the naturally-occurring amino acid substitutions in mammalian Robo3 (we refer to this variant as “Robo3NPLP”); second, we replaced *Drosophila* Robo3’s entire Ig1 domain with the Ig1 domain from mouse Robo3 (Robo3mR3Ig1); third, we replaced *Drosophila* Robo3’s Ig1 domain with the Ig1 domain from the *Drosophila*Frazzled (Fra) protein, which is unrelated to the Robo family and does not bind Slit (Robo3FraIg1) (Kolodziej et al., 1996; Rajagopalan et al., 2000b). We tested each of these Robo3 variants in our Slit binding assay and found that all three receptor variants were stably expressed at the surface of cultured *Drosophila* cells. This indicates that none of these Ig1 modifications disrupted translation, membrane trafficking, or stability of the Robo3 protein. Cells expressing Robo3NPLP displayed strongly reduced, but still detectable, levels of Slit binding **(Figure 1D,I**). However, Slit did not bind to cultured cells expressing the Robo3mR3Ig1 or Robo3FraIg1 variants (similar to untransfected cells or cells expressing Robo3ΔIg1) **(Figure 1 E-F,I**). We next set out to characterize the axonal localization and axon guidance activities of these Slit binding-deficient Ig1 variants *in vivo*.

### Slit binding-deficient Ig1 variants of Robo3 are properly localized *in vivo*

We used the same CRISPR strategy to replace the endogenous *Drosophila robo3* gene with each of the three variants described above (creating *Robo3*^*Robo3NPLP*^, *Robo3*^*Robo3mR3Ig1*^, and *Robo3*^*Robo3FraIg1*^alleles), and examined the expression and localization of the HA-tagged variant proteins when expressed from the *Robo3* locus. We found that the expression of Robo3NPLP closely reproduced that of full-length Robo3: the majority of the protein was localized to longitudinal axons with relatively little detectable within neuronal cell bodies and was excluded from axons in the medial one-third of the neuropile **(Figure 5E)**. Robo3mR3Ig1 and Robo3FraIg1 were also primarily localized to axons, with only slightly elevated staining in neuronal cell bodies compared to full-length Robo3 **(Figure 5F,G**). These results confirm that interaction with Slit is not required for the Ig1-dependent axonal localization of Robo3. This suggests that whatever residues or structural characteristics are required for Ig1’s role in Robo3 localization must be at least partially conserved in the Ig1 domain of mouse Robo3 and *Drosophila*Fra.

We observed that axons expressing Robo3NPLP or Robo3mR3Ig1 in place of Robo3 are properly excluded from the medial zone, suggesting that both of these variants can rescue Robo3’s ability to guide axons into the intermediate and lateral zones **(Figure 5E,F**; Figure 6G-J). In contrast, HA-positive axons were not properly excluded from the medial zone in homozygous *Robo3*^*Robo3FraIg1*^ embryos, suggesting that this activity is at least partially compromised in the Robo3FraIg1 variant **(Figure 5G)**. Consistent with this, exclusion of Robo3FraIg1-expressing axons from the medial zone was restored in heterozygous *Robo3*^*Robo3FraIg1*^*/+*animals as described for Robo3ΔIg1 and Robo1ΔIg1 above **(Figure 6K,L**).

### Slit binding-deficient Ig1 variants of Robo3 fully rescue longitudinal pathway formation

Next, we examined FasII-positive longitudinal pathways in *Robo3*^*Robo3NPLP*^, *Robo3*^*Robo3mR3Ig1*^, and *Robo3*^*Robo3FraIg1*^ embryos to quantify the extent to which these variants can substitute for *Robo3*to guide intermediate longitudinal axons. We found that intermediate FasII pathways formed properly in 98.6% and 97.2% of hemisegments in homozygous *Robo3*^*Robo3NPLP*^and *Robo3*^*Robo3mR3Ig1*^ embryos, respectively, confirming that both Robo3NPLP and Robo3mR3Ig1 can fully substitute for Robo3 to promote formation of these axon pathways **(Figure 2J,K**). This, combined with the observation that Robo3mR3Ig1 does not exhibit detectable Slit binding, suggests that Robo3 guides longitudinal axons to the intermediate zone independently of Slit. Notably, FasII-positive longitudinal pathways formed correctly in only 62.9% of hemisegments in homozygous *Robo3*^*Robo3FraIg1*^ embryos (a similar degree of rescue to *Robo3*^*Robo3ΔIg1*^), despite Robo3FraIg1’s relatively normal axonal localization **(Figure 2L)**. This partial rescue suggests that the Robo3FraIg1 variant lacks an Ig1-dependent guidance function that is present in both Robo3NPLP and Robo3mR3Ig1 and genetically separable from its role in axonal localization. Thus, it appears that Robo3 Ig1 contributes to longitudinal axon guidance through a function that is independent of Slit binding and separable from its role in axonal localization, and which is shared by Robo3NPLP and Robo3mR3Ig1, but not Robo3FraIg1.

## Discussion

Here we have examined the functional importance of the Ig1 domain of the *Drosophila* Robo3 receptor for Slit binding, *in vivo*protein localization, and axon guidance in the *Drosophila* embryonic ventral nerve cord. Deleting the Ig1 domain from Robo3 prevents its interaction with Slit in cultured cells and interferes with the axonal localization of Robo3 protein in embryonic neurons. However, the Robo3ΔIg1 protein can partially substitute for full-length Robo3 in promoting the formation of longitudinal axon pathways in the intermediate region of the nerve cord. Additional Robo3 variants in which the Ig1 domain has been modified or replaced completely also exhibit reduced or absent Slit binding ability, yet completely rescue Robo3-dependent intermediate pathway formation, indicating that this axon guidance activity of Robo3 is independent of its ability to bind Slit.

### *Drosophila* Robo3’s N-terminal Ig1 domain is required for Slit binding

Most Robo family receptors interact with their canonical Slit ligands via an evolutionarily conserved binding interface involving the Ig1 domain of the Robo proteins and the LRR2 (D2) domain of Slit ligands (Fukuhara et al., 2008; Liu et al., 2004; Morlot et al., 2007). There is one *slit* gene in *Drosophila*, and the Slit protein it encodes has been shown to bind to all three *Drosophila* Robo receptors (Robo1, Robo2, and Robo3) (Brose et al., 1999; Rajagopalan et al., 2000b). We have previously shown that the Ig1 domains of *Drosophila* Robo1 and Robo2 are required for Slit binding (Brown et al., 2015; Evans et al., 2015), while the other ectodomain elements in Robo1 are dispensable for Slit binding (Brown and Evans, 2020; Brown et al., 2018; Reichert et al., 2016).

Here we show that a version of Robo3 from which the Ig1 domain has been deleted (Robo3ΔIg1) does not display detectable Slit binding in cultured *Drosophila* cells. Thus, like its paralogs Robo1 and Robo2, the Ig1 domain of *Drosophila* Robo3 is also necessary for interaction with Slit.

### Slit-dependent vs Slit-independent functions of Robo3 Ig1

The experiments described in this paper have distinguished three genetically separable functions of Robo3’s Ig1 domain: 1) Slit binding, 2) axonal trafficking/localization, and 3) guidance of longitudinal axons. Robo3 Ig1 is necessary for Slit binding, since Robo3ΔIg1 displays no detectable interaction with Slit in our ligand binding assays. Robo3 Ig1 is also partially required for axonal localization and intermediate pathway formation, as these are both partially compromised when endogenous Robo3 is replaced by Robo3ΔIg1. Robo3mR3Ig1 and Robo3FraIg1 are both primarily localized to axons *in vivo*, but neither displays detectable Slit binding, indicating that the Slit binding function of Ig1 can be separated from its role in axonal localization. The axonal localization function of Ig1 is also separable from its activity in longitudinal axon guidance, since Robo3mR3Ig1 and Robo3FraIg1 exhibit similar axonal localization but Robo3mR3Ig1 completely rescues intermediate pathway formation while Robo3FraIg1 only rescues as well as Robo3ΔIg1, which is not properly localized. Thus it appears that Slit binding via Ig1 is not required for proper axonal localization of Robo3, or for its axon guidance role in longitudinal pathway formation. Ig1-dependent localization of Robo3 may depend on specific Ig1 sequences that are shared in *Drosophila* Robo3, mouse Robo3, and *Drosophila*Fra, or instead it may depend on a structural conformation provided by five tandem Ig domains rather than any particular Ig1 sequence. Structural studies of wild type and variant ectodomains and/or more restricted sequence swaps may provide further insight into this question.

The axon guidance defects in *Robo3*^*Robo3ΔIg1*^ embryos cannot be accounted for by lack of Slit binding, since Robo3mR3Ig1 has no detectable Slit binding yet fully rescues longitudinal axon guidance. They also cannot be accounted for by decreased axonal localization of Robo3ΔIg1, since Robo3FraIg1 is axonally localized yet only rescues to a similar extent as Robo3ΔIg1. Thus Robo3 Ig1 must have some function that is independent of Slit binding, and independent of its role in axonal localization, in guiding intermediate axons. This activity is not disrupted by the NPLP double mutation and is shared by *Drosophila* Robo1 Ig1 and mouse Robo3 Ig1 (since Robo3NPLP, Robo1, and Robo3mR3Ig1 all fully rescue Robo3-dependent intermediate pathway formation), but not *Drosophila* Fra Ig1 (since Robo3FraIg1 guides intermediate axons only as well as Robo3ΔIg1).

### Additional domains or sequences outside of Ig1 contribute to Robo3’s axon guidance function

Robo3ΔIg1 can partially substitute for full-length Robo3 to guide intermediate longitudinal axons, indicating that additional sequences or domains outside of Ig1 also contribute to its axon guidance activity. This secondary contribution is not conserved in *Drosophila* Robo1 (since *Robo3*^*robo1ΔIg1*^ does not rescue intermediate pathway defects) and does not appear to be necessary when a Robo family Ig1 domain is present (since *Robo3*^*robo1*^ fully rescues intermediate pathway defects). This secondary non-Ig1 guidance function cannot substitute for loss of Ig1 even when axonal localization is restored, because Robo3FraIg1 is properly localized and should retain any non-Ig1 guidance function(s), but does not completely rescue intermediate pathway defects.

It should be possible to identify which domain(s) contribute to this secondary function(s) by deleting other ectodomain elements, either alone or in combination with Ig1. We are currently using the CRISPR gene replacement approach described here to assay the functional importance of the remaining Ig and Fn domains of Robo3 via systematic individual domain deletions, similar to our investigations of Robo1 structural domains (Brown and Evans, 2020; Brown et al., 2015; Brown et al., 2018; Reichert et al., 2016). A summary of the activities present in Robo1 and Robo3 variants and how they relate to functional properties of Ig1 and/or other regions of the receptors is presented in **Table 2**.

**Table 2.**
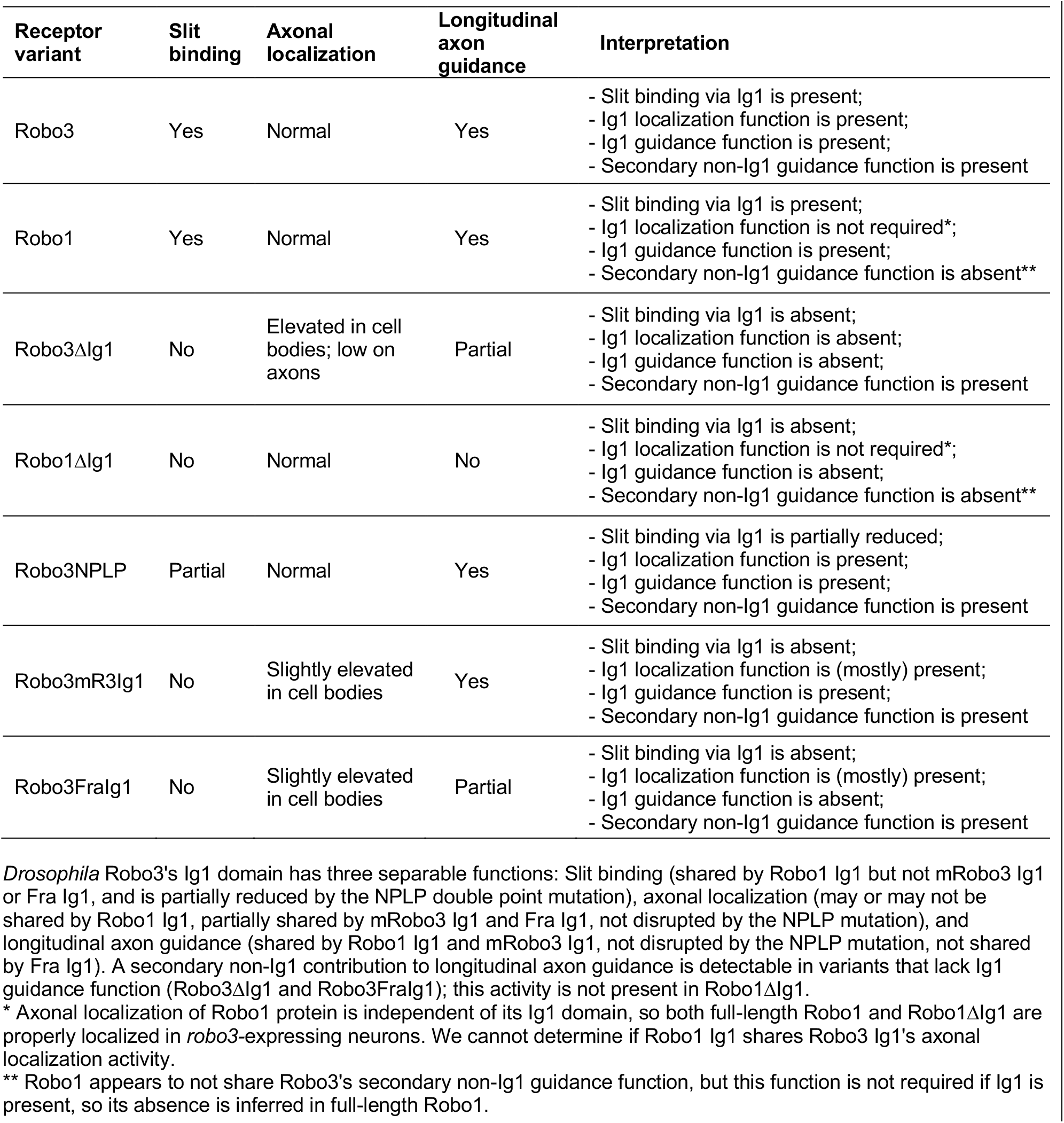
Summary of activities present in Robo3 and Robo1 variants

### Differential regulation of Robo receptor trafficking and localization

Our results show that Robo1ΔIg1 protein is localized correctly in Robo1-expressing and Robo3-expressing neurons, while Robo3ΔIg1 is mislocalized in both populations of neurons. We have previously described a similar mislocalization of Robo2ΔIg1 protein in Robo2-expressing neurons (a relative increase in cell body protein levels and decrease in axonal protein levels) (Howard et al., 2021). These observations suggest that Ig1 domains may be playing different roles in protein trafficking and/or localization in Robo1 vs Robo2/Robo3, and that this difference is specific to the proteins themselves, not the population of neurons in which they are expressed.

We note that the increased cell body localization of Robo3ΔIg1 in Robo1-expressing neurons (which is strongly punctate or “speckly” throughout the cortex, where neuronal cell bodies are located, and individual cell body outlines are not readily apparent) appears qualitatively distinct from its localization in Robo3-expressing neurons (where the outlines of individual cell bodies can be easily distinguished) **(compare Figure 5C with Figure S1D)**. This may reflect differences in Robo protein trafficking in different neuronal subsets. The increased cell body localization we have previously reported for Robo2ΔIg1 appears similar to Robo3ΔIg1 localization, suggesting similarities in trafficking between Robo2 and Robo3 in their respective neuronal subsets (Howard et al., 2021).

The elevated punctate cell body localization of Robo3ΔIg1 in Robo1-expressing neurons resembles an effect that we have previously described for Robo1ΔIg3 and Robo1ΔFn1, although these Robo1 variants did not display the same degree of reduction in axonal localization as Robo3ΔIg1 **(Figure S1D-F**)(Brown et al., 2018; Reichert et al., 2016). The midline repulsive function of Robo1ΔIg3 and Robo1ΔFn1 is not compromised by this shift in localization, as both can fully substitute for full-length Robo1 to promote normal levels of midline repulsion (Brown et al., 2018; Reichert et al., 2016).

### Cell-autonomous vs non-autonomous axon guidance functions of *Drosophila* Robo2 and Robo3

*Drosophila* Robo2 and Robo3 are both required for the formation of longitudinal axon pathways in the embryonic ventral nerve cord. In the absence of *robo2*, lateral axon pathways fail to form correctly, while in the absence of *Robo3*, intermediate pathways fail to form correctly. The overlapping expression patterns of the three Robos initially suggested a model where they might act equivalently to confer quantitatively increasing levels of Slit repulsion based on how many Robo receptors an individual axon would express (Rajagopalan et al., 2000b; Simpson et al., 2000b), but later studies showed that their functions were not interchangeable (Evans and Bashaw, 2010; Spitzweck et al., 2010). Notably, Robo2 can substitute for Robo3 to guide intermediate axons, but Robo3 cannot substitute for Robo2 to guide lateral axons, suggesting that Robo2 might guide lateral axons via a distinct mechanism that is not shared by Robo3 (Spitzweck et al., 2010).

We have recently reported that Robo2ΔIg1-expressing axons are not misguided into the intermediate or medial zones in *robo2*^*robo2ΔIg1*^ embryos, even though FasII-positive lateral axon pathways fail to form correctly in these embryos. This suggests that Robo2 may guide FasII-positive lateral axons non-autonomously (Howard et al., 2021). Here we observe that Robo3-expressing axons are misguided into the medial zone in *Robo3*^*Robo3ΔIg1*^ embryos, suggesting that Robo3 guides longitudinal axons via a mechanism that is cell-autonomous and further distinguishing its likely mechanism of action from that of Robo2.

### Are Robo-dependent mechanisms of longitudinal axon guidance evolutionarily conserved?

Mammalian Robo receptors (Robo1 and Robo2) also regulate the guidance of longitudinal axons in the CNS, including medial-lateral sorting of post-crossing spinal commissural axons and longitudinal guidance of ipsilateral pioneer axons in the midbrain and hindbrain (Dugan et al., 2011; Farmer et al., 2008; Jaworski et al., 2010; Kim et al., 2014; Long et al., 2004; Mastick et al., 2010). In the case of mid- and hindbrain longitudinal tracts, guidance defects seen in *Robo1-/-; Robo2-/-*double mutant mice are reproduced in *Slit1-/-; Slit2-/-*double mutants, suggesting that Robos may guide pioneer axons in response to Slits (Farmer et al., 2008). However, while the loss of Robo1 and Robo2 results in meandering pioneer axons and axon tract disorganization, there appears to be no consistent directional shift in pioneer axon positions in *Robo1-/-*and *Robo2-/-*mutants, in contrast to the specific medial shifting defects in FasII pathways in fly *robo2* and *Robo3* mutants.

In the case of post-crossing spinal commissural axons, Robo1 and Robo2 appear to have opposing roles in guiding axons to ventral and lateral funiculi in mouse, as loss of Robo1 causes lateral shifting of axons, while loss of Robo2 causes ventral/medial shifting (Jaworski et al., 2010; Long et al., 2004). It is not clear whether post-crossing guidance of spinal commissural neurons by Robo1 and Robo2 is Slit-dependent.

In non-mammalian vertebrates, lateral positioning of longitudinal axon tracts in the zebrafish hindbrain is also regulated by Robo receptors, as longitudinal axons shift closer to the midline in *robo2/astray* mutants (Kastenhuber et al., 2009), and gain of function studies suggest that Robos can also influence longitudinal axon guidance in the chick spinal cord (Reeber et al., 2008). The roles of vertebrate Robos in longitudinal axon guidance have been interpreted as striking a balance between Slit-dependent repulsion and Netrin-dependent attraction to regulate medial-lateral position, and/or as promoting Slit-dependent fasciculation to sort longitudinal axons into specific pathways (Farmer et al., 2008; Kastenhuber et al., 2009; Kim et al., 2014). Our results support a Slit-independent function of *Drosophila* Robo3 in regulating intermediate pathway formation, suggesting that the molecular mechanisms underlying Robo-dependent longitudinal axon guidance may be distinct in insects and vertebrates.

## Materials and methods

### Molecular Biology

#### Robo3 and Robo1 variants

Robo3 domain deletion, point mutation and chimeric receptor variants created for this study were generated by PCR and include the following amino acids after the N-terminal 4×HA epitope tag (numbers refer to Genbank/NCBI reference sequences AAF51387 [*Drosophila* Robo3 = DmRobo3], NP_001158239 [mouse Robo3 = MmRobo3]), and AAF58493 [*Drosophila* Fra]): Robo3ΔIg1 (DmRobo3 L119-V1432), Robo3NPLP (DmRobo3 H21-V1432 with N43P and L82P point mutations), Robo3mR3Ig1 (MmRobo3 P64-V163 / DmRobo3 L119-V1432), Robo3FraIg1 (Fra L35-R142 / DmRobo3 L119-V1432). The Robo1ΔIg1 variant was described previously (Brown et al., 2015).

#### pUAST cloning

Robo3 coding sequences were cloned as BglII fragments into p10UASTattB for S2R+ cell transfection. All p10UASTattB constructs include identical heterologous 5′ UTR and signal sequences (derived from the *Drosophila wingless* gene) and an N-terminal 3×HA tag.

#### robo1 rescue construct cloning

Full-length Robo3 and Robo3ΔIg1 coding sequences were cloned as BglII fragments into the BamHI-digested *robo1* rescue construct backbone (Brown et al., 2015). Robo1 and Robo3 proteins produced from this construct include the endogenous Robo1 signal peptide and an N-terminal 4xHA tag. Coding regions were sequenced completely prior to injection.

#### Robo3 CRISPR donor and gRNA plasmid cloning

Construction of the initial *Robo3* CRISPR donor and gRNA plasmid was described previously (Evans, 2017). Full-length *Robo3* and *Robo3* variant cDNAs were cloned as BglII fragments into the rescue construct backbone, which includes an N-terminal 4xHA tag inserted in-frame after the first 5 bp of *Robo3* exon 2. Coding regions were sequenced completely prior to injection.

### Genetics

#### Drosophila strains

The following *Drosophila melanogaster* strains, mutant alleles, and transgenes were used: Canton-S (wild type), *Robo3*^*1*^(Rajagopalan et al., 2000b), *slit*^*2*^(also known as *slit*^*IG107*^)(Nüsslein-Volhard et al., 1984), *Robo3*^*Robo3*^(Spitzweck et al., 2010), *Robo3*^*robo1*^(Spitzweck et al., 2010), *Robo3*^*Robo3ΔIg1*^, *Robo3*^*robo1ΔIg1*^, *Robo3*^*Δ*^, *Robo3*^*Robo3NPLP*^, *Robo3*^*Robo3mR3Ig1*^, *Robo3*^*Robo3FraIg1*^, *P{robo1::HArobo1} (Brown et al*., *2015), p{robo1::HArobo3}, p{robo1::HArobo3ΔIg1}, w*^*1118*^; *sna*^*Sco*^*/CyO,P{en1}wg*^*en11*^*(“Sco/CyOwg”)*. Transgenic flies were generated by BestGene Inc (Chino Hills, CA) using ΦC31-directed site-specific integration into an attP landing site at cytological position 28E7 (for *robo1* genomic rescue constructs). All crosses were carried out at 25°C.

#### Generation and recovery of CRISPR modified alleles

The *Robo3*gRNA plasmid was co-injected with each homologous donor plasmid (*Robo3*^*Robo3ΔIg1*^, *Robo3*^*robo1ΔIg1*^, *Robo3*^*Robo3NPLP*^, *Robo3*^*Robo3mR3Ig1*^, *Robo3*^*Robo3FraIg1*^)into *nos-Cas9*.*P* embryos (Port et al., 2014) by BestGene Inc (Chino Hills, CA). Injected individuals (G0) were crossed as adults to *Sco/CyOwg*. Founders (G0 flies producing F1 progeny carrying modified *Robo3*alleles) were identified by testing two pools of three F1 females per G0 cross by genomic PCR with primers that produce diagnostic PCR product sizes when the expected modified allele is present. From each identified founder, 5-10 F1 males were then crossed individually to *Sco/CyOwg* virgin females. After three days, the F1 males were removed from the crosses and tested by PCR to determine if they carried the modified allele. The *Robo3*^*Δ*^ NHEJ allele was identified by screening F1 males from a positive founder (injected with the *Robo3*^*Robo3ΔIg1*^ donor) using PCR primers 589 and 590, which produce a smaller (approx. 2 kb) product when the NHEJ deletion is present. F2 flies from positive F1 crosses were used to generate balanced stocks, and the modified alleles were fully sequenced by amplifying the entire modified locus (approx. 6 kb) from genomic DNA using primers 589 and 590, then sequencing the PCR product after cloning via CloneJET PCR cloning kit (Thermo Scientific). Details of G0 survival, fertility, and modified allele transmission rates are provided in **Table 1**.

### Immunofluorescence and imaging

*Drosophila* embryo collection, fixation, and antibody staining were carried out as previously described (Hauptman et al., 2022; Patel, 1994). The following antibodies were used: FITC-conjugated goat anti-HRP (Jackson ImmunoResearch #123-095-021, 1:100), mouse anti-Fasciclin II (Developmental Studies Hybridoma Bank [DSHB] #1D4, 1:100), mouse anti-βgal (DSHB #40-1a, 1:150), mouse anti-Robo3 cytoplasmic (DSHB #15H2, 1:100), mouse anti-HA (Covance #MMS-101P-500, 1:1000), Cy3-conjugated goat anti-mouse (Jackson #115-165-003, 1:1000). Embryos were genotyped using balancer chromosomes carrying lacZ markers. Ventral nerve cords from embryos of the desired genotype and developmental stage were dissected and mounted in 70% glycerol/PBS. Fluorescent confocal stacks were collected using a Leica SP5 confocal microscope and processed by Fiji/ImageJ (Schindelin et al., 2012) and Adobe Photoshop software.

### Slit binding assay

*Drosophila*S2R+ cells were cultured at 25ºC in Schneider’s media plus 10% fetal calf serum. To assay Slit binding, cells were plated on uncoated 12 mm circular glass coverslips in six-well plates (Robo-expressing cells) or 75 cm^2^ cell culture flasks (Slit-expressing cells) at a density of 1-2×10^6^ cells/ml and transfected with pRmHA3-GAL4 (Klueg et al., 2002) and HA-tagged p10UAST-Robo or untagged pUAST-Slit plasmids using Effectene transfection reagent (Qiagen). GAL4 expression was induced with 0.5 mM CuSO_4_ for 24 hours, then Slit-conditioned media was harvested by adding heparin (2.5 ug/ml) to Slit-transfected cells and incubating at room temperature for 20 minutes with gentle agitation. Robo-transfected cells were incubated with Slit-conditioned media at room temperature for 20 minutes, then washed with PBS and fixed for 20 minutes at 4ºC in 4% formaldehyde. Cells were permeabilized with PBS+0.1% Triton X-100, then stained with antibodies diluted in PBS+2mg/ml BSA. Antibodies used were: mouse anti-SlitC (Developmental Studies Hybridoma Bank [DSHB] #c555.6D, 1:50), rabbit anti-HA (Covance #PRB-101C-500, 1:2000), Cy3-conjugated goat anti-mouse (Jackson #115-165-003, 1:500), and Alexa 488-conjugated goat anti-rabbit (Jackson #111-545-003, 1:500). After antibody staining, coverslips with cells attached were mounted in Aqua-Poly/Mount (Polysciences, Inc.). Fluorescent confocal stacks were collected using a Leica SP5 confocal microscope and processed by Fiji/ImageJ (Schindelin et al., 2012) and Adobe Photoshop software. For quantification of Slit binding, confocal max projections were opened in Fiji/ImageJ and individual cells were outlined using the “Freehand selections” tool. Pixel intensities for the anti-Slit and anti-HA (Robo) channels were measured for 14-24 cells for each Robo3 variant, and the average ratio of Slit:HA for each construct (normalized to full-length Robo3) is reported as “relative Slit binding” in Fig 1.

## Acknowledgments

Stocks obtained from the Bloomington Drosophila Stock Center [National Institutes of Health (NIH) grant P40 OD-018537) were used in this study. Monoclonal antibodies were obtained from the Developmental Studies Hybridoma Bank, created by the Eunice Kennedy Shriver National Institute of Child Health and Human Development of the NIH and maintained at The Department of Biology, University of Iowa, Iowa City, IA 52242. This work was supported by NIH grant R15 NS-098406 (T.A.E.).

## Supplemental Figure S1

**Figure S1.**
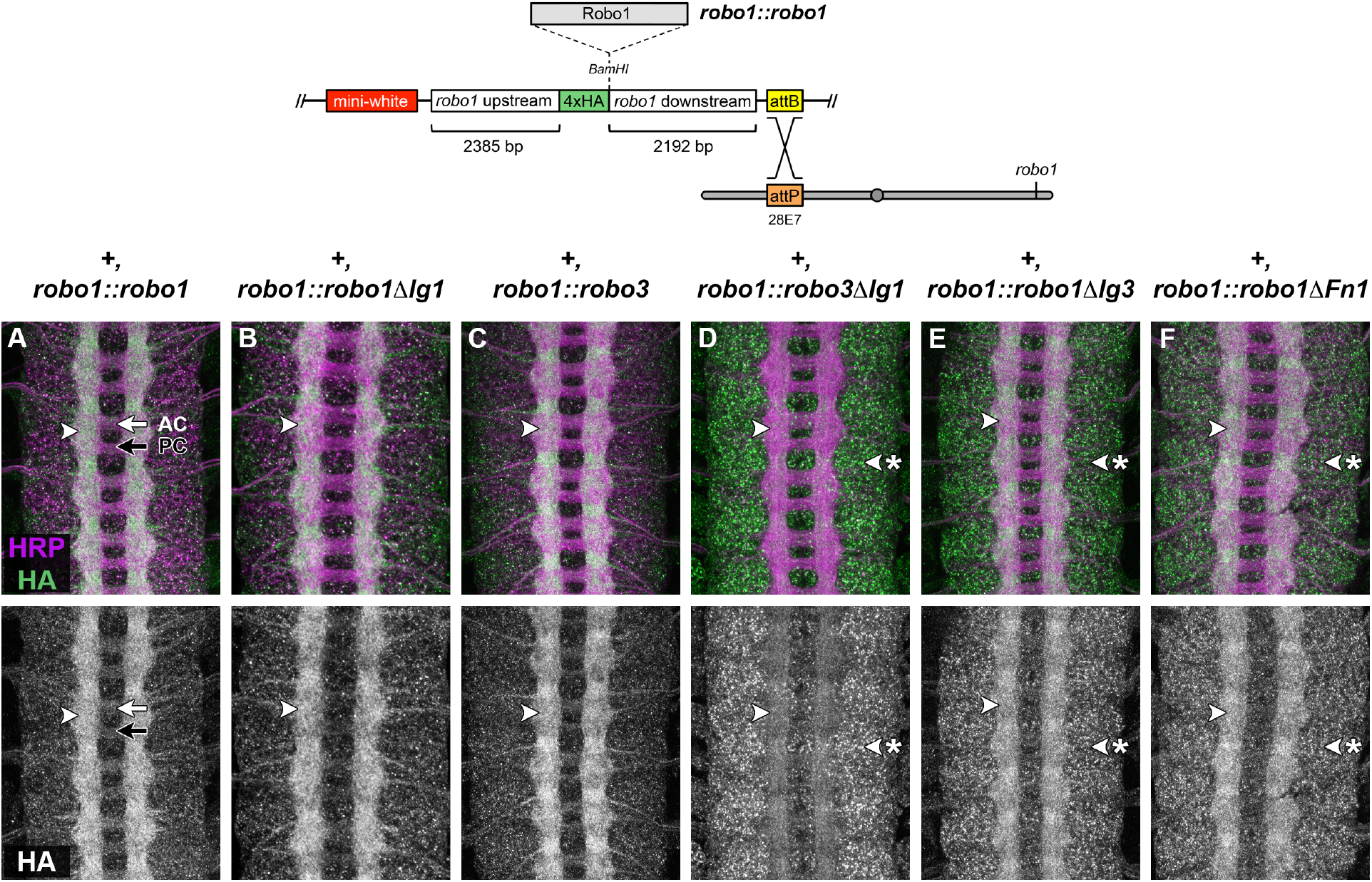
Expression of Robo1 and Robo3 variants in *robo1-*expressing neurons. **(A-E)**Stage 16-17 *Drosophila*embryonic ventral nerve cords stained with anti-HRP (magenta; labels all axons) and anti-HA (green) antibodies. Lower images show anti-HA channel alone from the same embryos. All embryos carry a *robo1*rescue transgene driving expression of the indicated variant in the neurons that normally express *robo1*, in an otherwise wild type background. HA-tagged full-length Robo1 **(A)**Robo1ΔIg1 **(B)**, or full-length Robo3 **(C)**proteins expressed from the *robo1* rescue transgene are localized to longitudinal axons **(arrowhead)** and are excluded from commissural axon segments in both the anterior commissure **(AC, white arrow)**and posterior commissure **(PC, black arrow)**. The HA staining pattern in these embryos closely matches the expression of endogenous Robo1 protein. Robo3ΔIg1 protein expressed from the same transgene exhibits lower expression on axons **(D, arrowhead)**and elevated expression in neuronal cell bodies **(D, arrowhead with asterisk)**. We have previously observed similarly elevated cell body localization of Robo1ΔIg3 and Robo1ΔFn1 proteins, although both of these Robo1 variants also retain high-level axonal localization, unlike Robo3ΔIg1 **(E**,**F)**(Brown et al., 2018; Reichert et al., 2016). Schematic of the *robo1* rescue construct is illustrated above (Brown et al., 2015).

